# Differential Regulation of Hepatic Akt/mTOR Signaling During Acute and Chronic *Toxoplasma gondii* Infection in a Murine Model

**DOI:** 10.64898/2026.04.06.716682

**Authors:** Jianchun Xiao

## Abstract

*Toxoplasma gondii* is an obligate intracellular parasite that infects virtually all warm-blooded animals, progressing through acute and chronic stages. The Akt/mTOR signaling axis plays critical roles in cell survival, proliferation, and metabolism, making it a key target for intracellular pathogens. This study investigated how *T. gondii* infection modulates this pathway during both infections. Outbred CD-1 mice were infected intraperitoneally with the virulent GT1 strain of *T. gondii*. Mice for acute studies were sacrificed five days post-infection, while those for chronic studies were treated with sulfadiazine and sacrificed five months post-infection. Phosphoprotein expression of eight Akt/mTOR pathway components was measured in liver tissues using a multiplexed bead-based immunoassay. Acute *T. gondii* infection caused broad suppression of Akt/mTOR signaling, with 6 of 8 markers significantly downregulated, including pS6RP^Ser235/236^, pAKT^S473^, pBAD^Ser136^, pIRS1^S636/639^, pPTEN^Ser380^, and pGSK-3α/β^Ser21/9^. In contrast, chronic infection selectively activates specific nodes of the pathway in a cyst burden-dependent manner, including pBAD^Ser136^, pmTOR^Ser2448^, and pGSK-3α/β^Ser21/^9. There are strong correlations in signaling changes between inter-components, which reflect coherent and coordinated pathway-level reprogramming rather than random perturbation. These findings show that acute and chronic *T. gondii* infections have opposing effects on host Akt/mTOR signaling for their own benefit, which may present new therapeutic targets.

**Graphical Abstract:** 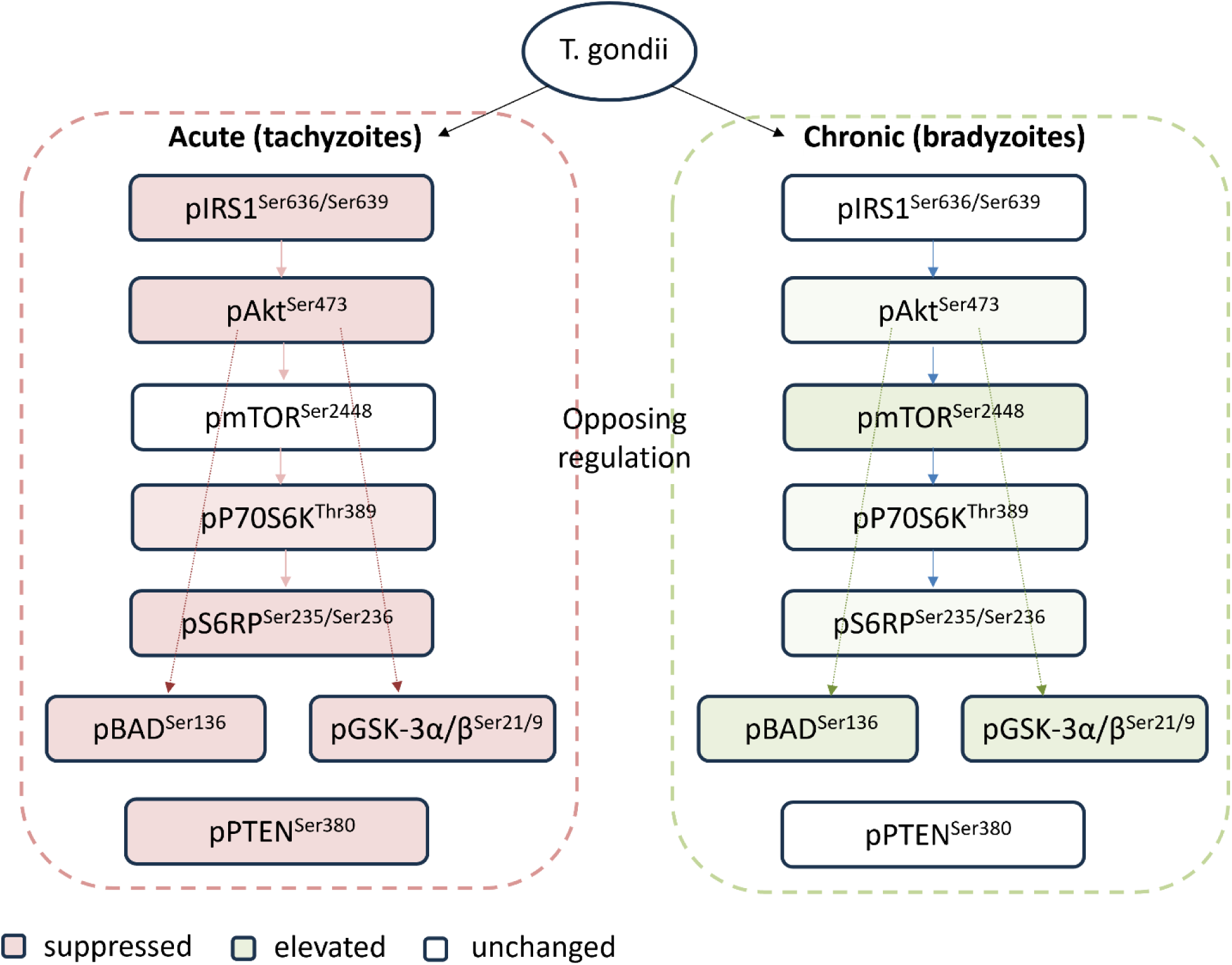

**Highlights:**

- Acute *T. gondii* infection broadly suppresses hepatic Akt/mTOR signaling
- Chronic infection exerts cyst burden-dependent activation of specific Akt/mTOR nodes
- *T. gondii* has distinct strategies to manipulate host survival based on its life stages.
- The Akt/mTOR pathway may serve as a therapeutic target for the treatment of *T. gondii*.

## Introduction

*Toxoplasma gondii* is an obligate intracellular parasite that infects approximately one-third of the global human population, making it one of the most prevalent parasitic infections worldwide. The infection develops in two stages: acute and chronic [1]. During an acute infection, rapidly dividing tachyzoites disseminate throughout the host, prompting strong innate and adaptive immune responses to control the multiplication of the parasite. This immune response leads to the conversion of tachyzoites into the slower-growing encysted bradyzoites. Tissue cysts can persist for the lifetime of the host and remain infectious if ingested. Currently, available anti-*T. gondii* drugs are ineffective in clearing the cyst form of the chronic infection [2]. This therapeutic gap largely results from an incomplete understanding of interactions between the host and the parasite during the progression of *T. gondii* infection. Gaining insight into how *T. gondii* infection affects cell survival, metabolism, and proliferation during acute and chronic infection may lead to better approaches.

The ability to subvert host cell signaling pathways is central to intracellular parasite survival. The phosphatidylinositol 3-kinase (PI3K)/Akt/mammalian target of rapamycin (mTOR) pathway is a master regulator of cell survival, metabolism, and growth [3]. Upon activation by growth factors or insulin receptor signaling, PI3K phosphorylates phosphatidylinositol-4,5-bisphosphate (PIP2) to generate PIP3. This recruits Akt to the plasma membrane, where it is phosphorylated at Thr308 by PDK1 and at Ser473 by the mTOR complex 2 (mTORC2) [4]. Fully activated Akt coordinates diverse downstream responses, including: inhibition of apoptosis through phosphorylation of the pro-apoptotic protein BAD at Ser136, which makes BAD dissociate from the Bcl-2/Bcl-X complex and lose the pro-apoptotic function [5]; inhibition of glycogen synthase kinase-3α/β (GSK-3α/β) through phosphorylation at Ser21/9, promoting cell survival and glucose metabolism [6]; and activation of mTOR complex 1 (mTORC1) through multiple mechanisms, leading to phosphorylation of downstream effectors including p70 ribosomal S6 kinase 1 (p70S6K1) at Thr389 and ribosomal protein S6 (S6RP), which drive ribosome biogenesis and protein synthesis. Negative regulation of the pathway is mediated by the tumor suppressor PTEN, which dephosphorylates PIP3 to dampen Akt activation. Additionally, an inhibitory phosphorylation of insulin receptor substrate-1 (IRS1) at Ser636/639, a negative feedback mechanism downstream of mTORC1.

Research has suggested that several intracellular pathogens have evolved mechanisms to manipulate the PI3K/Akt/mTOR pathway to their advantage. Mycobacterium tuberculosis [7], Salmonella [8], and Listeria [9] exploit Akt signaling to inhibit apoptosis and promote intracellular survival. The PI3K/Akt/mTOR signaling plays a very important role in HPV-induced carcinogenesis by regulating multiple cellular and molecular events [10]. Prior work has demonstrated that *T. gondii* can activate PI3K/Akt/mTOR signaling to facilitate host cell invasion and inhibit apoptosis in infected cells [11,12]. A recent study suggests that PI3K/Akt/mTOR signaling pathway plays an important role in *T. gondii*-induced mitochondrial dysfunction and the reprogramming of cellular energy metabolism [13]. However, systematic profiling of the complete Akt/mTOR signaling network across both acute and chronic infection stages has not been reported.

The current study systematically profiled the phosphorylation of eight components of the Akt/mTOR pathway in mouse liver tissues, during both acute and chronic *T. gondii* infection, using a multiplexed immunoassay. The hypothesis was that *T. gondii* modulates hepatic Akt/mTOR signaling differently based on its tachyzoite and bradyzoite life stages. The liver is a key metabolic organ and an important site of *T. gondii* infection. During the early stages of infection, tachyzoites disseminate via the bloodstream and infect hepatocytes [14,15]. While significant attention has focused on central nervous system toxoplasmosis, the liver can also harbor bradyzoite cysts during chronic infection [16]. Research has found that the liver is the primary site of tissue pathology [14] in severe toxoplasmosis and can cause liver diseases such as hepatitis [17] and hepatomegaly [18,19]. The present study revealed that tachyzoites rapidly replicate within a metabolically suppressed but still-viable host cell, while bradyzoites require a stable, long-lived cellular niche for decades-long persistence.

## Materials and Methods

### Ethics statement

All mouse specimens were collected from prior projects, and no additional animals were sacrificed for the present study. The animals were humanely euthanized in accordance with the Animal Protection Protocols at Johns Hopkins University and following the National Institutes of Health Guide for the Care and Use of Laboratory Animals.

### Acute and chronic mouse models of *T. gondii* infection

Six- to eight-week-old female outbred CD-1 mice (ICR-Harlan Sprague) were infected intraperitoneally (i.p.) with 500 *T. gondii* GT1 strain tachyzoites (Type I, virulent). Control mice received vehicle only (PBS). The model employs a type I strain because of its close association with clinical disease and its greater influence on host genes, as demonstrated in our previous research [20,21]. For acute infection, mice were sacrificed at 5 days post infection (dpi), as described previously [22]. For chronic infection, infected mice, including controls, were treated with anti-*T. gondii* chemotherapy (sulfadiazine sodium) in drinking water (400 mg/liter; Sigma) from days 5 to 30 to control tachyzoite proliferation and prevent animal death, and were sacrificed at five months postinfection (mpi) [23]. Upon sacrifice, liver tissue was immediately harvested, snap-frozen in liquid nitrogen, and stored at −80°C.

### Confirmation of infection and cyst burden assessment

As described previously [23], *T. gondii* infection in all chronically infected mice was confirmed using a commercial ELISA kit (IB19213, IBL America) for anti-*T. gondii* IgG antibodies. Cyst burden was quantified using the MAG1 (matrix antigen 1) assay, which measures antibodies to peptide antigens derived from MAG1 [24,25]. The MAG1 protein is an antigen found abundantly in the cyst wall and within the cyst matrix. The serological response to the MAG1 peptide antigen has been validated as a proxy marker of chronic *T. gondii* infection and cyst burden [23]. The median MAG1 antibody absorbance value (0.5) was the cutoff to stratify mice into two groups: those with a high cyst burden (OD ≥ 0.5) and those with a low cyst burden (OD < 0.5).

### Liver tissue homogenization and protein extraction

Liver tissue samples (approximately 100 mg) were homogenized on ice in RIPA buffer (Sigma), supplemented with a protease and phosphatase inhibitor cocktail (Thermo Scientific). The mixture was sonicated at 4°C for 5 min, followed by centrifugation at 10,000 × g for 5 minutes at 4°C, and the supernatant was collected. Total protein concentration was determined using a BCA protein assay (Thermo Scientific).

### Multiplexed Phosphoprotein Immunoassay

Phosphorylation levels of Akt/mTOR pathway components were measured using a bead-based multiplex immunoassay (Bio-Plex Pro Cell Signaling Akt Panel 8-plex, LQ00006JK0K0RR, Bio-Rad) on the Luminex xMAP platform. The panel comprises eight key phosphoproteins located upstream and downstream from Akt and includes: insulin receptor substrate-1 at Ser636/639 (pIRS1^Ser636/Ser639^), phosphatase and tensin homolog at Ser380 (pPTEN^Ser380^), serine/threonine-protein kinase Akt-1 at Ser 473 (pAkt^Ser473^), glycogen synthase kinase-3α/β at Ser21 and Ser9 (pGSK3α/β^Ser21/Ser9^), mTOR at Ser2448 (pmTOR^Ser2448^), p70 S6 kinase at Thr389 (pP70S6K^Thr389^), ribosomal protein S6 kinase beta-1 at Ser235/236 (pS6RP^Ser235/Ser236^), and the Bcl2-associated agonist of cell death at Ser136 (pBAD^Ser136^). All samples were analyzed according to the manufacturer’s protocols, and the median fluorescence intensity (MFI) values were recorded.

### Statistical Analysis

All data sets were tested for normality by using the Shapiro-Wilk test. Differences between two groups were analyzed by Welch’s t-test. For multiple groups, ANOVA with Bonferroni’s multiple comparison test was used. Spearman correlation analyses were performed to examine the inter-protein correlation. Fold change (FC) was calculated as the ratio of the mean MFI of the infection group to the mean MFI of the respective control group. A p-value of < 0.05 was considered statistically significant; p < 0.10 was noted as a trend (†). Data are presented as mean ± standard deviation (SD). All statistical analyses were conducted in Graph-Pad Prism V11.0.0.

## Results

### Acute *T. gondii* infection broadly suppresses hepatic Akt/mTOR signaling

To examine the effects of acute *T. gondii* infection on host Akt/mTOR signaling, the phosphorylation levels of eight pathway components in liver tissues was compared between mock-infected and acutely infected mice at 5 dpi.

As shown in Table 1, acute infection resulted in significant suppression of Akt/mTOR signaling across most examined nodes. The most pronounced reductions were observed for p70S6K^Thr389^ (FC = 0.616, p = 0.056), pIRS1^Ser636/Ser639^ (FC = 0.636, p = 0.017), pS6RP^Ser235/Ser236^ (FC = 0.630, p = 0.016), and pAKT^Ser473^ (FC = 0.694, p = 0.013), with each showing a 31–38% decrease compared to controls. A moderate decrease was also observed for pBAD^Ser136^ (FC = 0.753, p = 0.010), pGSK-3α/β^Ser21/9^ (FC = 0.783, p = 0.011), and pPTEN^Ser380^ (FC = 0.868, p = 0.033). Notably, pmTOR^Ser2448^ showed a minimal reduction (FC = 0.862) and did not reach statistical significance (p = 0.181). Overall, 6 of 8 markers were significantly downregulated in infected mice, with p70S6K^Thr389^ showing a trend towards reduction.

**Table 1.**
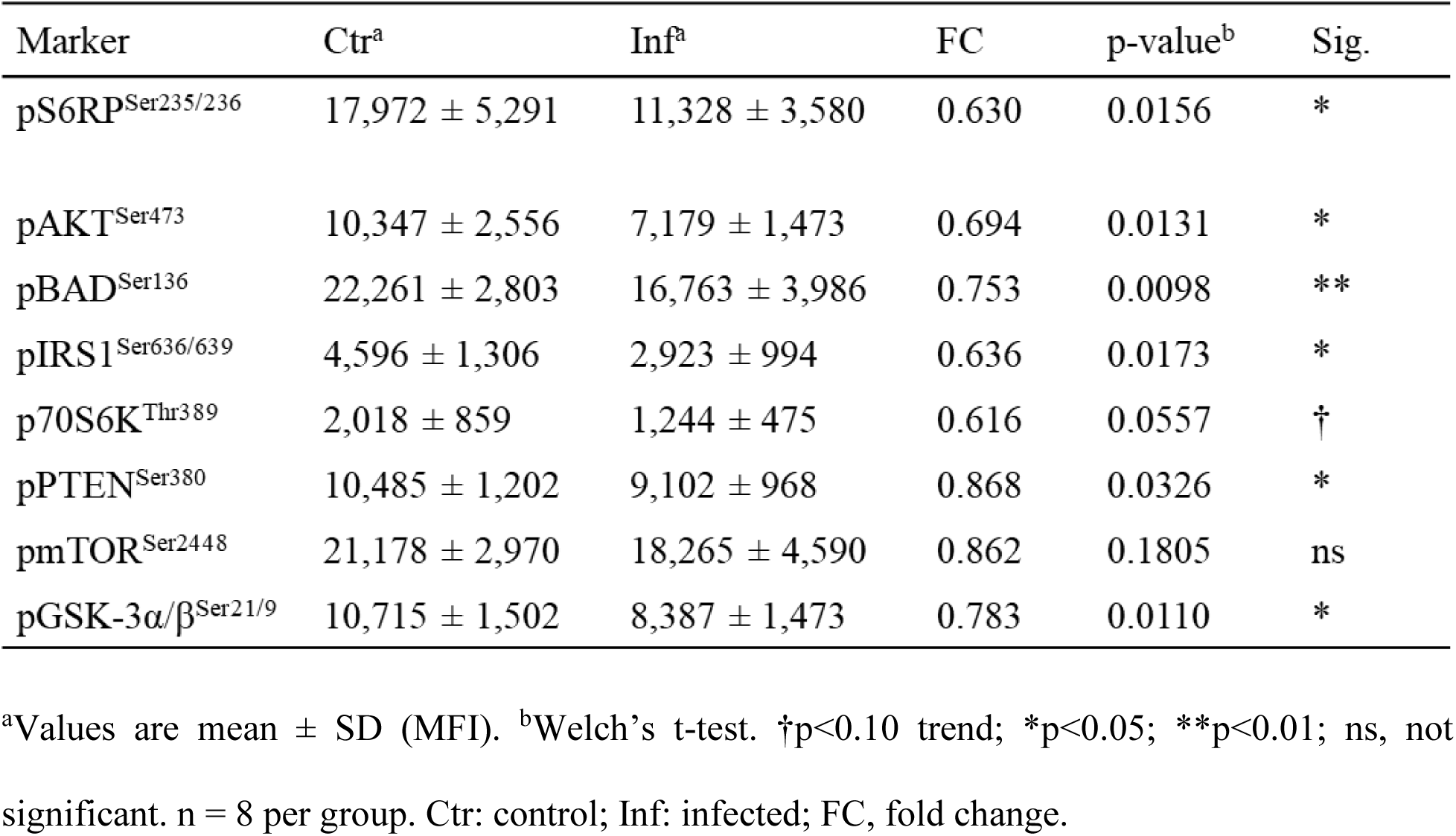
Hepatic Akt/mTOR phosphoprotein levels during acute *T. gondii* infection.

### Chronic *T. gondii* infection elicits cyst burden-dependent activation of select hepatic Akt/mTOR nodes

Chronic *T. gondii* infection activated specific nodes of the Akt/mTOR signaling pathway in a cyst burden-dependent manner (Table 2). Compared to mock-infected control, mice that were chronically infected for 5 months showed increased phosphorylation of all 8 components, with significant changes in four: pGSK-3α/β^Ser21/9^ (FC = 1.72, p < 0.0001), pmTOR^Ser2448^ (FC = 1.44, p = 0.0118), pBAD^Ser136^ (FC = 1.82, p = 0.0082), and pAKT^Ser473^ (FC = 1.41, p = 0.0077). Additionally, p70S6K^Thr389^ showing a trend towards increase (FC = 1.92, p = 0.0607).

**Table 2.**
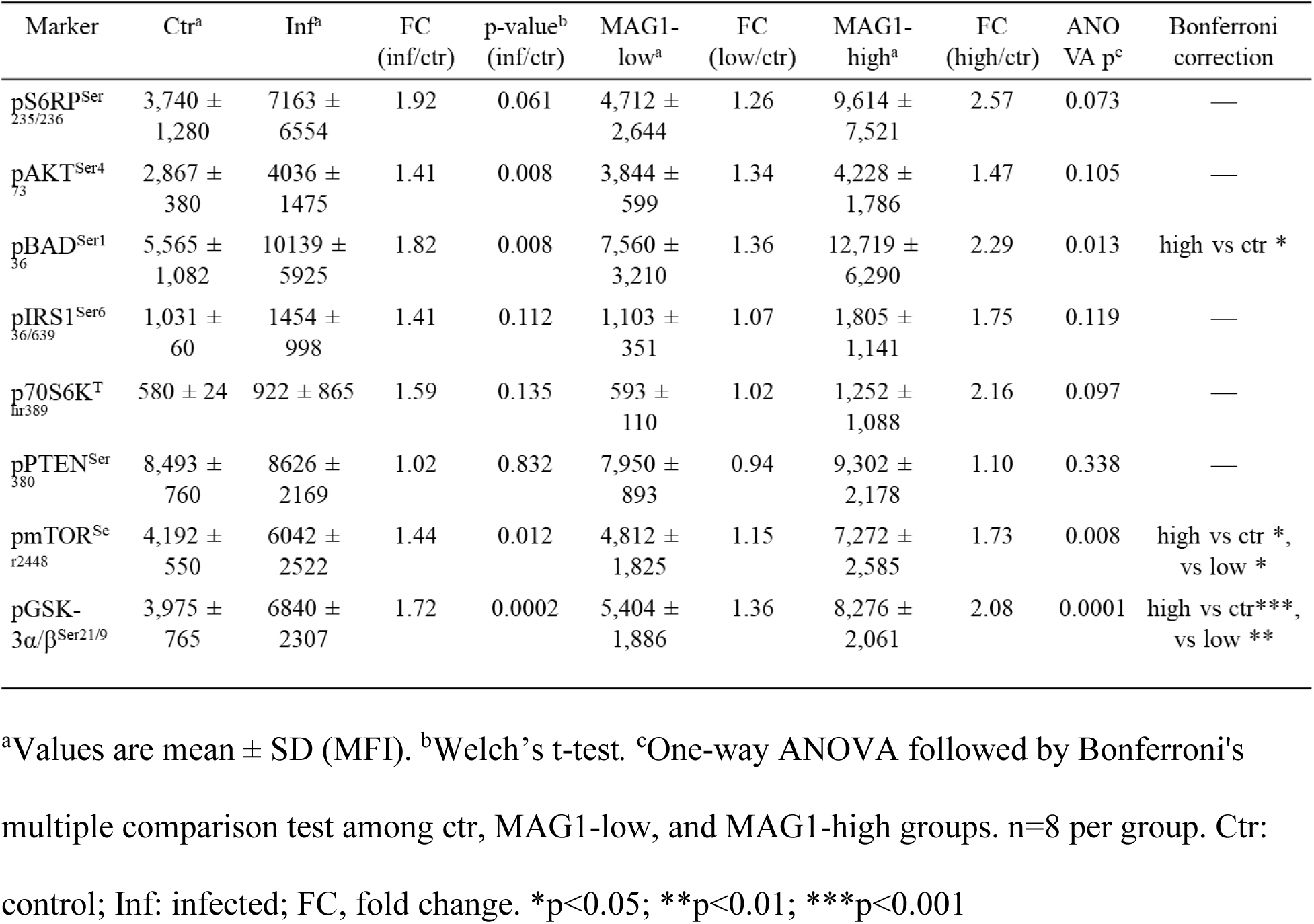
Hepatic Akt/mTOR phosphoprotein levels during chronic *T. gondii* infection.

Chronic infection is defined by the presence of tissue cysts, making cyst burden a critical factor for understanding its effects. I stratified the chronically infected mice into MAG1-high and MAG1-low groups based on MAG1 antibody levels, which serve as a proxy marker of cyst burden [23]. The results indicated that the elevation was particularly high in MAG1-high mice; 5 of 8 pathway components showed a significant or near-significant increase compared to controls (Table 2). Specifically, pGSK-3α/β^Ser21/9^ showed a greater than twofold increase (FC = 2.08, p < 0.0001), as did pBAD^Ser136^ (FC = 2.29, p = 0.013). Additionally, pmTOR^Ser2448^ was elevated (FC = 1.73, p = 0.010), while p70S6K^Thr389^ (FC = 2.16, ANOVA p = 0.0972) and pS6RP^Ser235/Ser236^ (FC = 2.57, ANOVA p=0.0732) showed a trend towards increase. Moreover, MAG1-high mice also significantly differed from MAG1-low mice in pmTOR^Ser2448^ (FC = 1.51, p = 0.046) and pGSK-3α/β^Ser21/9^ (FC = 1.53, p = 0.004). No significant difference was found between MAG1-low and mock-infected controls.

None of the eight Akt/mTOR pathway components correlate significantly with MAG1 antibody levels in the chronically infected group (all p > 0.05 by Spearman). However, there is a trend indicating a positive correlation between MAG1 antibody levels and pGSK-3α/β^Ser21/9^ (r = 0.46, p = 0.0758) or pBAD^Ser136^ (r = 0.43, p = 0.096).

### Phosphorylation levels of Akt/mTOR pathway components are significantly correlated

I explored whether activation/inhibition of these Akt/mTOR components was consistent along the pathway. To this aim, correlation analyses were performed among the 8 proteins from both acute and chronic groups, as well as their corresponding controls (Fig. 1).

**Figure 1.**
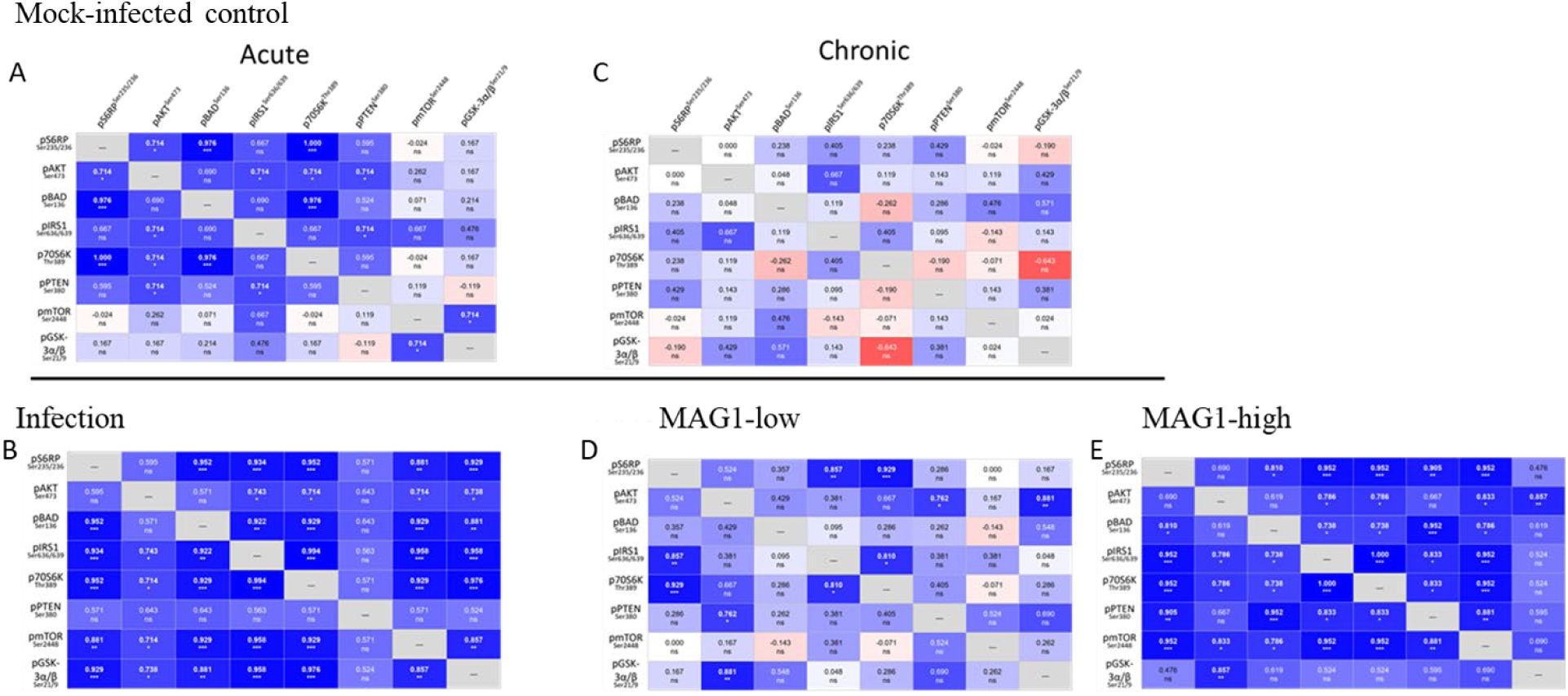
Correlation analyses in mock-infected controls as well as in acutely and chronically infected mice. Spearman correlation analyses were performed separately for each group: mock-infected controls during acute infection (A), acutely infected mice (B), mock-infected controls during chronic infection (C), MAG1-low mice (D), and MAG1-high mice (E) to examine associations among the components measured in the Akt/mTOR pathway. The color scheme of blue, white, and red was utilized to represent the strength and direction of the correlation coefficient. Blue: positive correlation; White: no correlation; Red: negative correlation. bold text (p < 0.05).

Mice with acute infections or high MAG1 antibody levels show a cohesive phosphoprotein network, with 19 statistically significant pairwise correlations. The weakest correlations during acute infection mainly involved pPTEN^Ser380^, whereas in mice with high MAG1 levels, the weakest correlations were primarily with pGSK-3α/β^Ser21/9^. The correlation between p70S6K^Thr389^ and pS6RP^Ser235/Ser236^ is the most robustly coupled link in every infection group (acute, MAG1-low, MAG1-high), confirming tight coupling between the kinase and its substrate during infection. For MAG1-low mice, pathway coherence is partially maintained but notably weaker, with only 5 pairwise correlations being significant. In the mock-infected control for chronic infection, all 28 pairwise correlations are non-significant, while 19 are non-significant in the mock-infected control for acute infection. The correlation between pAKT^Ser473^ and pBAD^Ser136^ remains non-significant across all groups, regardless of the infection status.

## Discussion

This study provides the first systematic, comparative analysis of hepatic Akt/mTOR signaling during both acute and chronic *T. gondii* infection in a murine model using the virulent GT1 strain. It revealed that distinct infection stages exert opposing regulatory effects on the host Akt/mTOR pathway: acute infection broadly suppresses activity across most nodes, whereas chronic infection selectively activates anti-apoptotic and metabolic signaling nodes in a parasite burden-dependent manner. These findings are consistent with the distinct functional imperatives of the tachyzoite and bradyzoite life stages [26,27], and support the concept that *T. gondii* has evolved stage-specific strategies to manipulate host cell biology for its own benefit.

### Broad suppression of Akt/mTOR signaling during acute infection

The broad suppression of hepatic Akt/mTOR signaling during acute infection — with 6 of 8 markers significantly reduced at 5 dpi — is consistent with metabolic exploitation and immune evasion during the lytic tachyzoite cycle [28,29].

Upstream of Akt, the reduction in pIRS1^Ser636/639^ appears insufficient to restore downstream Akt signaling, as concurrent activation of pPTEN^Ser380^ occurs. PTEN is a widely known negative regulator of insulin/PI3K signaling [30], so its reduction at Ser380 renders PTEN more enzymatically active, leading to further dephosphorylation of PIP3 and reinforcing suppression of the PI3K/Akt axis. Phosphorylation of IRS1 at Ser636/639 normally results from mTORC1/S6K1-mediated negative feedback on insulin signaling [31]; its reduction most likely reflects the loss of this feedback loop, as supported by reduced p70S6K^Thr389^ and pS6RP^Ser235/236^ expression. This reduction would be expected to enhance IRS1 activity and insulin sensitivity [32]. Pro-inflammatory cytokines (TNF-α, IL-1β, IL-6) activated during acute toxoplasmosis are well-known to interfere with insulin receptor signaling [33].

Downstream of Akt, reduced phosphorylation of BAD^Ser136^ increases pro-apoptotic pressure on the host cell by allowing BAD to interact with the anti-apoptotic proteins Bcl-2 and Bcl-xL [5]. Suppression of Akt signaling may attenuate NF-κB-dependent inflammatory gene expression, since Akt can phosphorylate and activate IκB kinase [34]. Dampening pro-inflammatory responses could thereby delay the onset of effective Th1 immunity. The phosphorylation of GSK3β^Ser9^ or GSK3α^Ser21^ is known to decrease GSK3α/β enzymatic activity [35,36]. The decrease in pGSK-3α/β indicates GSK-3 activation, which reflects a shift away from anabolic glucose storage and pro-survival signaling. The suppression of p70S6K^Thr389^ and pS6RP^Ser235/236^ may redirect biosynthetic resources away from host cell metabolism [37]. This would favor nutrient availability for the rapidly replicating tachyzoite within the parasitophorous vacuole.

Notably, mTOR phosphorylation at Ser2448 was the only node that did not show a significant change during acute infection. This implies that mTOR catalytic activity is preserved through Akt-independent mechanisms [38]. This preservation is biologically critical because mTOR is the primary suppressor of autophagy, and its activity prevents autophagic targeting of the parasitophorous vacuole — the host cell’s principal mechanism for intracellular parasite clearance [39,40]. Collectively, this pattern suggests that *T. gondii* has evolved to uncouple mTOR from its upstream Akt activator and converted the hepatocyte into a nutrient-rich but immunologically crippled replication niche.

### Selective activation of Akt/mTOR nodes during chronic infection

In contrast to acute infection, chronic *T. gondii* infection selectively activates specific nodes of anti-apoptotic (pBAD), metabolic (pmTOR, pGSK-3α/β), and translational (p-70S6K, p-S6RP) pathways in a parasite burden-dependent manner. This pattern suggests an overactivated signaling state, supporting the notion that bradyzoites inhibit host cell apoptosis and modify glucose metabolism to create a favorable environment for their survival [26,27]. It is conceivable that MAG1-low mice have insufficient levels of antigens needed to activate these nodes.

The cyst burden-associated activation of pBAD^Ser136^ and pGSK-3α/β^Ser21/Ser9^ carries broad functional implications. Phosphorylation of BAD at Ser136 would enhance anti-apoptotic signaling and maintain host cell viability [41]. Previous studies have found that *T. gondii*-infected cells are broadly resistant to multiple inducers of apoptosis [42]. This resistance may result from Akt and BAD phosphorylation, which, depending on parasite load, inhibit Bax translocation to mitochondria and block apoptosis [12]. Inactivation of GSK-3 may enhance glycolysis and increase glucose availability [43], which is crucial for bradyzoites. GRA18, a dense granule effector protein of *T. gondii*, has been shown to act as an inhibitor of host GSK3 triggering β-catenin accumulation [44]. Beta-catenin is a key molecule in the Wnt pathway, and its stable expression on Treg cells promotes cell survival [45]. Notably, our previous study found that the phosphorylation levels of the inhibitory forms of GSK-3α/β^Ser21/Ser9^ were decreased in the cerebellum of MAG1-high mice [46], whereas in this study, a more than two-fold increase in the liver was observed. This tissue-specific divergence likely reflects the well-documented pleiotropy of GSK-3 across different cellular contexts.

The significant upregulation of Akt, mTOR, and mTOR-associated targets (p70S6K^Thr389^ and pS6RP^Ser235/236^) in chronically infected mice — particularly in the MAG1-high subgroup — aligns with previous findings of *T. gondii* infection on Akt/mTOR pathway activation [13,47]. Hyperactivation of mTOR has been associated with defects in autophagosome formation, and inhibiting mTOR activity suppresses *T. gondii* replication [48]. A study reports that *T. gondii* inhibits FOXO3a, a transcription factor that regulates the expression of autophagy-related genes, through AKT-dependent phosphorylation [49]. Interestingly, the upstream regulatory machinery of the Akt/mTOR pathway — pIRS1^Ser636/639^ and pPTEN^Ser380^ — was not significantly altered during chronic infection, despite robust changes in downstream nodes. The lack of change suggested that upstream PI3K/PTEN regulation may not be the primary driver of chronic-stage Akt activation. Rather, direct activation at or downstream of Akt itself may be responsible and warrants further investigation. Additionally, although cyst burden is a primary factor in the activation of the hepatic Akt/mTOR pathway, there was no significant correlation between phosphorylation levels and MAG1 antibody levels. This suggests that humoral immunity does not directly reflect the intracellular signaling status within the tissue.

### Opposing Stage-Specific Strategies

The most striking finding of this study is the diametrically opposed regulatory strategies employed by *T. gondii* between acute and chronic infections. Acute infection broadly suppresses this pathway to redirect host metabolism and dampen immune activation, while chronic infection selectively activates pro-survival, anti-apoptotic, and anabolic nodes in a parasite burden-dependent manner. Similarly, our previous studies also found that *T. gondii* affects host miR-132 differently in acute and chronic infection [22,50]. A study on neonatal mouse astrocytes found that *T. gondii* infection triggers both pro-apoptotic and anti-apoptotic signals [51]. The cells activate apoptosis signals shortly after infection; however, the parasite inhibits programmed cell death for up to 24 hours. This delay allows the parasite to replicate, egress, and ultimately cause cellular destruction.

There were highly coordinated changes in the phosphorylation levels of adjacent kinases within the Akt/mTOR pathway in infected groups. This indicates that the integrity of the signaling cascade is maintained in the liver and that the change is not a stochastic phenomenon but a coherent event at the pathway level. While pmTOR^Ser2448^ showed no change in acute infection, it has significant correlations with all other components except pPTEN^Ser380^. Given the correlation between pAKT^Ser473^ and pBAD^Ser136^ is consistently non-significant across all groups, this warrants caution in interpreting BAD as a direct readout of Akt activity. Notably, inter-component correlations within control groups differed between the acute and chronic experiments, with acute controls showing stronger pathway coherence for several pairs. This difference may reflect age-related changes in hepatic metabolism.

### Clinical and Pathological Implications

Previous clinical and epidemiological studies have established associations between *T. gondii* infection — including both acute and chronic toxoplasmosis — and chronic liver diseases, including hepatomegaly, granulomas, hepatitis, cirrhosis, and hepatocellular necrosis in both immunocompetent and immunocompromised patients [52–56]. These findings provide novel molecular mechanistic insight into these diseases, identifying the Akt/mTOR signaling axis as a stage-specific, dynamically regulated target of *T. gondii* infection.

The PI3K/Akt/mTOR pathway is also prominently dysregulated across a wide spectrum of human cancers [57,58]. Given that chronic toxoplasmosis can lead to prolonged overactivation of Akt/mTOR pathways, these findings suggest that long-term *T. gondii* infection could increase oncogenic risk, as noted in other studies [59,60].

One limitation of this study is the lack of correlations with conventional liver injury biomarkers, including alanine transaminase (ALT), aspartate aminotransferase (AST), total bilirubin, and alpha-fetoprotein (AFP). Integrating these clinical-biochemical parameters with the phosphoproteomic data reported here would strengthen the translational relevance of these findings.

## Conclusion

In conclusion, this study showed that *T. gondii* exerts stage-specific, opposing regulation of host hepatic Akt/mTOR signaling. These findings reveal the liver as an immunometabolically active site of parasite-host interaction across infection stages. These insights open new avenues for stage-specific therapeutic intervention against toxoplasmosis and provide a molecular framework for understanding the hepatopathological consequences of infection.

## Funding

This work was supported by the Stanley Medical Research Institute (SMRI).

## Data availability

Any additional information is available on request.

## Acknowledgments

The author thanks Dr Ye Li for technical support.

## Conflicts of Interest

The author declares that she has no competing interests.

## Notes

### Competing Interest Statement

The authors have declared no competing interest.

## References

1. Dubey, J.P. Advances in the life cycle of Toxoplasma gondii. Int J Parasitol 1998, 28, 1019–1024, doi:10.1016/s0020-7519(98)00023-x.

2. Konstantinovic, N.; Guegan, H.; Stajner, T.; Belaz, S.; Robert-Gangneux, F. Treatment of toxoplasmosis: Current options and future perspectives. Food Waterborne Parasitol 2019, 15, e00036, doi:10.1016/j.fawpar.2019.e00036.

3. Manning, B.D.; Toker, A. AKT/PKB Signaling: Navigating the Network. Cell 2017, 169, 381–405, doi:10.1016/j.cell.2017.04.001.

4. Hay, N.; Sonenberg, N. Upstream and downstream of mTOR. Genes Dev 2004, 18, 1926–1945, doi:10.1101/gad.1212704.

5. Laplante, M.; Sabatini, D.M. mTOR signaling in growth control and disease. Cell 2012, 149, 274–293, doi:10.1016/j.cell.2012.03.017.

6. Cantley, L.C. The phosphoinositide 3-kinase pathway. Science 2002, 296, 1655–1657, doi:10.1126/science.296.5573.1655.

7. Lachmandas, E.; Beigier-Bompadre, M.; Cheng, S.C.; Kumar, V.; van Laarhoven, A.; Wang, X.; Ammerdorffer, A.; Boutens, L.; de Jong, D.; Kanneganti, T.D.;, et al. Rewiring cellular metabolism via the AKT/mTOR pathway contributes to host defence against Mycobacterium tuberculosis in human and murine cells. Eur J Immunol 2016, 46, 2574–2586, doi:10.1002/eji.201546259.

8. Owen, K.A.; Meyer, C.B.; Bouton, A.H.; Casanova, J.E. Activation of focal adhesion kinase by Salmonella suppresses autophagy via an Akt/mTOR signaling pathway and promotes bacterial survival in macrophages. PLoS Pathog 2014, 10, e1004159, doi:10.1371/journal.ppat.1004159.

9. Gessain, G.; Tsai, Y.H.; Travier, L.; Bonazzi, M.; Grayo, S.; Cossart, P.; Charlier, C.; Disson, O.; Lecuit, M. PI3-kinase activation is critical for host barrier permissiveness to Listeria monocytogenes. J Exp Med 2015, 212, 165–183, doi:10.1084/jem.20141406.

10. Zhang, L.; Wu, J.; Ling, M.T.; Zhao, L.; Zhao, K.N. The role of the PI3K/Akt/mTOR signalling pathway in human cancers induced by infection with human papillomaviruses. Mol Cancer 2015, 14, 87, doi:10.1186/s12943-015-0361-x.

11. Seizova, S.; Ruparel, U.; Garnham, A.L.; Bader, S.M.; Uboldi, A.D.; Coffey, M.J.; Whitehead, L.W.; Rogers, K.L.; Tonkin, C.J. Transcriptional modification of host cells harboring Toxoplasma gondii bradyzoites prevents IFN gamma-mediated cell death. Cell Host Microbe 2022, 30, 232–247 e236, doi:10.1016/j.chom.2021.11.012.

12. Quan, J.H.; Cha, G.H.; Zhou, W.; Chu, J.Q.; Nishikawa, Y.; Lee, Y.H. Involvement of PI 3 kinase/Akt-dependent Bad phosphorylation in Toxoplasma gondii-mediated inhibition of host cell apoptosis. Exp Parasitol 2013, 133, 462–471, doi:10.1016/j.exppara.2013.01.005.

13. Xu, K.; Zhu, S.; Ma, J.; Zu, M.; Yang, J.; Xu, F.; Dai, L.; Liu, D.; Wang, Y.; Zhang, X.;, et al. PI3K/Akt/mTOR signaling pathway mediates energy metabolic reprogramming and regulates mitochondrial homeostasis in host cells exposed to Toxoplasma gondii. Microbiol Spectr 2026, 14, e0138525, doi:10.1128/spectrum.01385-25.

14. Mordue, D.G.; Monroy, F.; La Regina, M.; Dinarello, C.A.; Sibley, L.D. Acute toxoplasmosis leads to lethal overproduction of Th1 cytokines. J Immunol 2001, 167, 4574–4584, doi:10.4049/jimmunol.167.8.4574.

15. Ferguson, D.J. Use of molecular and ultrastructural markers to evaluate stage conversion of Toxoplasma gondii in both the intermediate and definitive host. Int J Parasitol 2004, 34, 347–360, doi:10.1016/j.ijpara.2003.11.024.

16. Skariah, S.; McIntyre, M.K.; Mordue, D.G. Toxoplasma gondii: determinants of tachyzoite to bradyzoite conversion. Parasitol Res 2010, 107, 253–260, doi:10.1007/s00436-010-1899-6.

17. Atilla, A.; Aydin, S.; Demirdoven, A.N.; Kilic, S.S. Severe Toxoplasmic Hepatitis in an Immunocompetent Patient. Jpn J Infect Dis 2015, 68, 407–409, doi:10.7883/yoken.JJID.2014.422.

18. Huang, J.; Zhang, H.; Liu, S.; Wang, M.; Wan, B.; Velani, B.; Zhu, Y.; Lin, S. Is Toxoplasma gondii infection correlated with nonalcoholic fatty liver disease?- a population-based study. BMC Infect Dis 2018, 18, 629, doi:10.1186/s12879-018-3547-1.

19. Babekir, A.; Mostafa, S.; Minor, R.C.; Williams, L.L.; Harrison, S.H.; Obeng-Gyasi, E. The Association of Toxoplasma gondii IgG and Liver Injury in US Adults. Int J Environ Res Public Health 2022, 19, doi:10.3390/ijerph19127515.

20. Xiao, J.; Buka, S.L.; Cannon, T.D.; Suzuki, Y.; Viscidi, R.P.; Torrey, E.F.; Yolken, R.H. Serological pattern consistent with infection with type I Toxoplasma gondii in mothers and risk of psychosis among adult offspring. Microbes Infect 2009, 11, 1011–1018, doi:10.1016/j.micinf.2009.07.007.

21. Xiao, J.; Jones-Brando, L.; Talbot, C.C., Jr.; Yolken, R.H. Differential effects of three canonical Toxoplasma strains on gene expression in human neuroepithelial cells. Infect Immun 2011, 79, 1363–1373, doi:10.1128/IAI.00947-10.

22. Xiao, J.; Li, Y.; Prandovszky, E.; Karuppagounder, S.S.; Talbot, C.C., Jr.; Dawson, V.L.; Dawson, T.M.; Yolken, R.H. MicroRNA-132 dysregulation in Toxoplasma gondii infection has implications for dopamine signaling pathway. Neuroscience 2014, 268, 128–138, doi:10.1016/j.neuroscience.2014.03.015.

23. Xiao, J.; Li, Y.; Prandovszky, E.; Kannan, G.; Viscidi, R.P.; Pletnikov, M.V.; Yolken, R.H. Behavioral Abnormalities in a Mouse Model of Chronic Toxoplasmosis Are Associated with MAG1 Antibody Levels and Cyst Burden. PLoS Negl Trop Dis 2016, 10, e0004674, doi:10.1371/journal.pntd.0004674.

24. Xiao, J.; Viscidi, R.P.; Kannan, G.; Pletnikov, M.V.; Li, Y.; Severance, E.G.; Yolken, R.H.; Delhaes, L. The Toxoplasma MAG1 peptides induce sex-based humoral immune response in mice and distinguish active from chronic human infection. Microbes Infect 2013, 15, 74–83, doi:10.1016/j.micinf.2012.10.016.

25. Xiao, J.; Bhondoekhan, F.; Seaberg, E.C.; Yang, O.; Stosor, V.; Margolick, J.B.; Yolken, R.H.; Viscidi, R.P. Serological Responses to Toxoplasma gondii and Matrix Antigen 1 Predict the Risk of Subsequent Toxoplasmic Encephalitis in People Living With Human Immunodeficiency Virus (HIV). Clin Infect Dis 2021, 73, e2270–e2277, doi:10.1093/cid/ciaa1917.

26. Cerutti, A.; Blanchard, N.; Besteiro, S. The Bradyzoite: A Key Developmental Stage for the Persistence and Pathogenesis of Toxoplasmosis. Pathogens 2020, 9, doi:10.3390/pathogens9030234.

27. Sullivan, W.J., Jr.; Jeffers, V. Mechanisms of Toxoplasma gondii persistence and latency. FEMS Microbiol Rev 2012, 36, 717–733, doi:10.1111/j.1574-6976.2011.00305.x.

28. Laliberte, J.; Carruthers, V.B. Host cell manipulation by the human pathogen Toxoplasma gondii. Cell Mol Life Sci 2008, 65, 1900–1915, doi:10.1007/s00018-008-7556-x.

29. Blader, I.J.; Saeij, J.P. Communication between Toxoplasma gondii and its host: impact on parasite growth, development, immune evasion, and virulence. APMIS 2009, 117, 458–476, doi:10.1111/j.1600-0463.2009.02453.x.

30. Chen, Z.; Dempsey, D.R.; Thomas, S.N.; Hayward, D.; Bolduc, D.M.; Cole, P.A. Molecular Features of Phosphatase and Tensin Homolog (PTEN) Regulation by C-terminal Phosphorylation. J Biol Chem 2016, 291, 14160–14169, doi:10.1074/jbc.M116.728980.

31. Copps, K.D.; White, M.F. Regulation of insulin sensitivity by serine/threonine phosphorylation of insulin receptor substrate proteins IRS1 and IRS2. Diabetologia 2012, 55, 2565–2582, doi:10.1007/s00125-012-2644-8.

32. Shehata, M.F. Role of the IRS-1 and/or -2 in the pathogenesis of insulin resistance in Dahl salt-sensitive (S) rats. Heart Int 2009, 4, e6, doi:10.4081/hi.2009.e6.

33. Popko, K.; Gorska, E.; Stelmaszczyk-Emmel, A.; Plywaczewski, R.; Stoklosa, A.; Gorecka, D.; Pyrzak, B.; Demkow, U. Proinflammatory cytokines Il-6 and TNF-alpha and the development of inflammation in obese subjects. Eur J Med Res 2010, 15 Suppl 2, 120–122, doi:10.1186/2047-783x-15-s2-120.

34. Song, Y.S.; Narasimhan, P.; Kim, G.S.; Jung, J.E.; Park, E.H.; Chan, P.H. The role of Akt signaling in oxidative stress mediates NF-kappaB activation in mild transient focal cerebral ischemia. J Cereb Blood Flow Metab 2008, 28, 1917–1926, doi:10.1038/jcbfm.2008.80.

35. Cross, D.A.; Alessi, D.R.; Cohen, P.; Andjelkovich, M.; Hemmings, B.A. Inhibition of glycogen synthase kinase-3 by insulin mediated by protein kinase B. Nature 1995, 378, 785–789, doi:10.1038/378785a0.

36. Peyrollier, K.; Hajduch, E.; Blair, A.S.; Hyde, R.; Hundal, H.S. L-leucine availability regulates phosphatidylinositol 3-kinase, p70 S6 kinase and glycogen synthase kinase-3 activity in L6 muscle cells: evidence for the involvement of the mammalian target of rapamycin (mTOR) pathway in the L-leucine-induced up-regulation of system A amino acid transport. Biochem J 2000, 350 Pt 2, 361–368.

37. Bahrami, B.F.; Ataie-Kachoie, P.; Pourgholami, M.H.; Morris, D.L. p70 Ribosomal protein S6 kinase (Rps6kb1): an update. J Clin Pathol 2014, 67, 1019–1025, doi:10.1136/jclinpath-2014-202560.

38. Bhatt, A.P.; Damania, B. AKTivation of PI3K/AKT/mTOR signaling pathway by KSHV. Front Immunol 2012, 3, 401, doi:10.3389/fimmu.2012.00401.

39. Jung, C.H.; Ro, S.H.; Cao, J.; Otto, N.M.; Kim, D.H. mTOR regulation of autophagy. FEBS Lett 2010, 584, 1287–1295, doi:10.1016/j.febslet.2010.01.017.

40. Cheng, A.; Zhang, H.; Chen, B.; Zheng, S.; Wang, H.; Shi, Y.; You, S.; Li, M.; Jiang, L. Modulation of autophagy as a therapeutic strategy for Toxoplasma gondii infection. Front Cell Infect Microbiol 2022, 12, 902428, doi:10.3389/fcimb.2022.902428.

41. Zha, J.; Harada, H.; Yang, E.; Jockel, J.; Korsmeyer, S.J. Serine phosphorylation of death agonist BAD in response to survival factor results in binding to 14-3-3 not BCL-X(L). Cell 1996, 87, 619–628, doi:10.1016/s0092-8674(00)81382-3.

42. Nash, P.B.; Purner, M.B.; Leon, R.P.; Clarke, P.; Duke, R.C.; Curiel, T.J. Toxoplasma gondii-infected cells are resistant to multiple inducers of apoptosis. J Immunol 1998, 160, 1824–1830.

43. Wang, L.; Li, J.; Di, L.J. Glycogen synthesis and beyond, a comprehensive review of GSK3 as a key regulator of metabolic pathways and a therapeutic target for treating metabolic diseases. Med Res Rev 2022, 42, 946–982, doi:10.1002/med.21867.

44. He, H.; Brenier-Pinchart, M.P.; Braun, L.; Kraut, A.; Touquet, B.; Coute, Y.; Tardieux, I.; Hakimi, M.A.; Bougdour, A. Characterization of a Toxoplasma effector uncovers an alternative GSK3/beta-catenin-regulatory pathway of inflammation. Elife 2018, 7, doi:10.7554/eLife.39887.

45. Ding, Y.; Shen, S.; Lino, A.C.; Curotto de Lafaille, M.A.; Lafaille, J.J. Beta-catenin stabilization extends regulatory T cell survival and induces anergy in nonregulatory T cells. Nat Med 2008, 14, 162–169, doi:10.1038/nm1707.

46. Xiao, J.; Huang, J.; Yolken, R.H. Elevated matrix Metalloproteinase-9 associated with reduced cerebellar perineuronal nets in female mice with toxoplasmosis. Brain Behav Immun Health 2024, 36, 100728, doi:10.1016/j.bbih.2024.100728.

47. Wang, Y.; Weiss, L.M.; Orlofsky, A. Intracellular parasitism with Toxoplasma gondii stimulates mammalian-target-of-rapamycin-dependent host cell growth despite impaired signalling to S6K1 and 4E-BP1. Cell Microbiol 2009, 11, 983–1000, doi:10.1111/j.1462-5822.2009.01305.x.

48. Leroux, L.P.; Lorent, J.; Graber, T.E.; Chaparro, V.; Masvidal, L.; Aguirre, M.; Fonseca, B.D.; van Kempen, L.C.; Alain, T.; Larsson, O.;, et al. The Protozoan Parasite Toxoplasma gondii Selectively Reprograms the Host Cell Translatome. Infect Immun 2018, 86, doi:10.1128/IAI.00244-18.

49. Diez, A.F.; Leroux, L.P.; Chagneau, S.; Plouffe, A.; Gold, M.; Chaparro, V.; Jaramillo, M. Toxoplasma gondii inhibits the expression of autophagy-related genes through AKT-dependent inactivation of the transcription factor FOXO3a. mBio 2023, 14, e0079523, doi:10.1128/mbio.00795-23.

50. Li, Y.E.; Kannan, G.; Pletnikov, M.V.; Yolken, R.H.; Xiao, J. Chronic infection of Toxoplasma gondii downregulates miR-132 expression in multiple brain regions in a sex-dependent manner. Parasitology 2015, 142, 623–632, doi:10.1017/S003118201400167X.

51. Contreras-Ochoa, C.O.; Lagunas-Martinez, A.; Belkind-Gerson, J.; Diaz-Chavez, J.; Correa, D. Toxoplasma gondii invasion and replication within neonate mouse astrocytes and changes in apoptosis related molecules. Exp Parasitol 2013, 134, 256–265, doi:10.1016/j.exppara.2013.03.010.

52. Ustun, S.; Aksoy, U.; Dagci, H.; Ersoz, G. Incidence of toxoplasmosis in patients with cirrhosis. World J Gastroenterol 2004, 10, 452–454, doi:10.3748/wjg.v10.i3.452.

53. Neves, E.S.; Bicudo, L.N.; Curi, A.L.; Carregal, E.; Bueno, W.F.; Ferreira, R.G.; Amendoeira, M.R.; Benchimol, E.; Fernandes, O. Acute acquired toxoplasmosis: clinical-laboratorial aspects and ophthalmologic evaluation in a cohort of immunocompetent patients. Mem Inst Oswaldo Cruz 2009, 104, 393–396, doi:10.1590/s0074-02762009000200039.

54. Hryzhak, I.H. Infection with Toxoplasma gondii can promote chronic liver diseases in HIV-infected individuals. Folia Parasitol 2020, 67, doi:10.14411/fp.2020.034.

55. Mohamed, B.M.; Omran, M.M.; Abdelrazek, M.A.; Attallah, A.M.; El-Far, M. Prevalence of Toxoplasma gondii 36-KDa antigen and chronic Hepatitis C: another evidence of an Association. J Parasit Dis 2021, 45, 1049–1054, doi:10.1007/s12639-021-01351-8.

56. El-Sayed, N.M.; Ramadan, M.E.; Ramadan, M.E. Toxoplasma gondii Infection and Chronic Liver Diseases: Evidence of an Association. Trop Med Infect Dis 2016, 1, doi:10.3390/tropicalmed1010007.

57. Janku, F.; Yap, T.A.; Meric-Bernstam, F. Targeting the PI3K pathway in cancer: are we making headway? Nat Rev Clin Oncol 2018, 15, 273–291, doi:10.1038/nrclinonc.2018.28.

58. Bossler, F.; Hoppe-Seyler, K.; Hoppe-Seyler, F. PI3K/AKT/mTOR Signaling Regulates the Virus/Host Cell Crosstalk in HPV-Positive Cervical Cancer Cells. Int J Mol Sci 2019, 20, doi:10.3390/ijms20092188.

59. Fadel, E.F.; Tolba, M.E.M.; Ahmed, A.M.; El-Hady, H.A. Serological and molecular detection of Toxoplasma Gondii among cancer patients in Sohag, Upper Egypt: a case-control study. Sci Rep 2025, 15, 5236, doi:10.1038/s41598-025-88680-3.

60. Mostafa, N.E.S.; Abdel Hamed, E.F.; Rashed, H.E.S.; Mohamed, S.Y.; Abdelgawad, M.S.; Elasbali, A.M. The relationship between toxoplasmosis and different types of human tumors. J Infect Dev Ctries 2018, 12, 137–141, doi:10.3855/jidc.9672.

